# A deep-learning-based screening platform in aging-relevant human motor neurons to identify therapeutic compounds for amyotrophic lateral sclerosis

**DOI:** 10.1101/2023.11.21.568079

**Authors:** Bhagawati Jalnapurkar, David Le, William Shainin, Ryan Zarcone, Rajat Bhatnagar, Joshua Chin, Conor Corbin, Charles Curt, Brian Hodge, Fernando Martinez, Thomas Mikita, Anne Pipathsouk, Jesse Rosen, Thomas A. Rando, Joseph T. Rodgers, Jeremy W. Linsley

## Abstract

Age is the primary risk factor for most neurodegenerative diseases including amyotrophic lateral sclerosis (ALS). Despite the clear importance of age on the development and progression of ALS, age is rarely considered a factor in cellular or animal models of ALS. The recent advent of direct reprogramming methodologies, whereby somatic cells such as fibroblasts can be converted into neurons while retaining the age signature of the host, has facilitated the study of age-relevant human cells *in vitro* for the first time. Despite the promise of this technology, age is a complex, multidimensional, and dynamic phenotype that makes the interpretation of the interplay between age and disease a great challenge. To circumvent these challenges, we developed a screening platform for directly reprogrammed neurons in an age-relevant human cellular model to better model and identify disease-modifying targets and pathways for ALS. This platform uses deep learning image analysis to screen compounds for efficacy against ALS-associated cellular phenotypes and efficiently classifies ALS-like phenotypic signatures as well as predicts the age of the original fibroblast donor. Notably, our work identified NCB-0846, a TNIK/MAP4K7 inhibitor, as a novel compound with the ability to revert ALS-like phenotypes. Moreover, we show that chronic dosing of NCB-0846 in the SOD1G93A mouse model of ALS significantly reduced serum neurofilament-L levels. Our findings establish the power of a platform that combines direct reprogramming of human cells and deep learning as a powerful tool for combatting aging and age-related diseases.

## Introduction

Amyotrophic lateral sclerosis (ALS) is a relentless neurodegenerative disorder marked by the progressive loss of motor neurons and subsequent paralysis. Age is the greatest risk factor for ALS^1^, with the majority of ALS cases presenting in late middle age^2^. Despite ongoing research, the intricate molecular pathways that link aging with ALS onset remain largely elusive^3^. The aging process thus represents a therapeutic target for ALS, and therapeutic strategies that reverse cellular aging could also potentially slow, halt, or even reverse ALS progression.

Given the strong age-dependence to the onset of ALS, there is a clear need for models that accurately reflect the age of neurons. Neurons derived from induced pluripotent stem cells (iPSC)s from patients with ALS lose aging signatures during reprogramming. Often these neurons do not reliably show ALS-relevant phenotypes, such as degeneration in culture, without the use of additional stressors or perturbations^4,5^. In contrast to iPSC-based methodologies, direct reprogramming allows for the conversion of a somatic cell, such as a fibroblast, into the cell type of interest without the cell trans-iting through a pluripotent, embryonic-like state. A number of direct reprogramming methodologies have been shown to preserve epi-genetic and functional aging signa-tures of donors^6,7^. By effectively retaining the age signature of the donor, directly reprogrammed neurons often show specific features of neurodegenerative disease that have been hard to recapitulate in iPSC-derived neurons. Such features include inclusion body formation in Huntington’s disease, seed competent Tau and mature Tau isoforms in tauopathies, Aß deposition in Alzheimer’s disease, and neuronal death in many pathologies^8–10^. These methodologies offer the promise of fully understanding the important interplay of age and neurodegenerative disease in human cells.

The onset and progression of ALS are complex and multifaceted. Adding the complex dynamics of aging exacerbates the challenge of disentangling the etiology of the disease. Machine learning techniques can leverage the scale of large datasets to identify trends in data that might be impossible to observe otherwise. Recently, the emergence of image-based deep learning (DL) algorithms has brought the use of machine learning to studies of biomedical research with powerful results, including the ability to identify signatures of neurodegenerative diseases such as ALS^11,12^ that were previously unattainable. Furthermore, application of DL in the analysis of cellular phenotypes often surpasses the ability of conventional analytical techniques to identify particular biological features or phenomena^13–15^. Deep learning’s application to microscopy is limited by the requirement for very large datasets to train these powerful models. The low reprogramming efficiency of direct reprogramming methodologies by viruses^16–20^ poses a particular challenge for generating large enough datasets of age-relevant directly reprogrammed cells for DL.

Here we use a chemical direct reprogramming platform and a methodology for small molecule screening of motor neurons to identify compounds capable of reverting age-relevant ALS phenotypes. We employ this platform to study neurons overexpressing TDP43^M337V^ or SOD1^G93A^, dominant negative mutations linked to familial ALS and used to model ALS^21–23^. This provides a high-resolution, multidimensional, phenotypic view of the disease within motor neurons. We use these models to identify compounds that reverse or blunt the effects of ALS-related phenotypes. When administered to a SOD1^G93A^ mouse model, a TNIK/MAP4K7 inhibitor identified from our *in vitro* screen shows efficacy at reducing neurofilament light chain (NfL) levels in serum. This powerful combination of direct reprogramming and DL analysis could expedite the discovery of novel, effective therapeutic strategies for ALS, serving as a roadmap for future studies in age-associated neurodegenerative disorders.

## Results

### A rapid screening platform for chemically reprogrammed motor neurons

Deep learning requires a large number of image examples to learn from in order to perform classification tasks. Achieving the scale and speed of data generation required for DL and small molecule screening has been particularly challenging for viral-based direct reprogramming methodologies, which often suffer from low reprogramming efficiencies that limit the usable biological examples per reprogramming cycle^18,24^. Additionally, many published direct reprogramming protocols require time courses of weeks to months for reprogramming^23^. These long timeframes increase the likelihood of cell heterogeneity, thus increasing assay noise due to batch effects. For these reasons, we looked to adapt a protocol in which reprogramming efficiency was high and the time course for reprogramming was as short as possible.

Adapting a recently described chemical reprogramming protocol for induced motor neurons (iMNs)^25^, we achieved a fast reprogramming time course for reprogramming that could be performed in 384-well plates with full liquid handling automation, facilitating acquisition of large datasets required for DL and HTS (Figure 1A-D). After the re-programming, converted iMNs showed increased anti-islet and anti-ChAT signal, markers of motor neuron fate, compared to unconverted fibroblasts (Figure 1B–D). Another advantage of chemical reprogramming protocols over viral reprogramming protocols is that virus can be used to express biosensors or transgenes in the iMNs without interference of other viruses used for reprogramming. We reasoned that lentiviral transduction of a neuron-specific biomarker in fibroblasts could be used as a selection marker for re-programmed neurons. Induced motor neurons showed strong lentiviral expression from a neuron-specific hSyn1 promoter^29^ compared to unconverted fibroblasts (Figure 1E– H). In a typical reprogramming cycle, we can process up to 32 plates, totaling over 1.4 million single iMN images per screening cycle, easily providing an adequate data set size for DL analysis. With the use of our lentiviral transduction labeling approach, iMN reprogramming could be imaged longitudinally, indicating iMNs could be labeled, identified, and genetically manipulated within live cultures in high throughput (Figure 1I).

**Figure 1.**
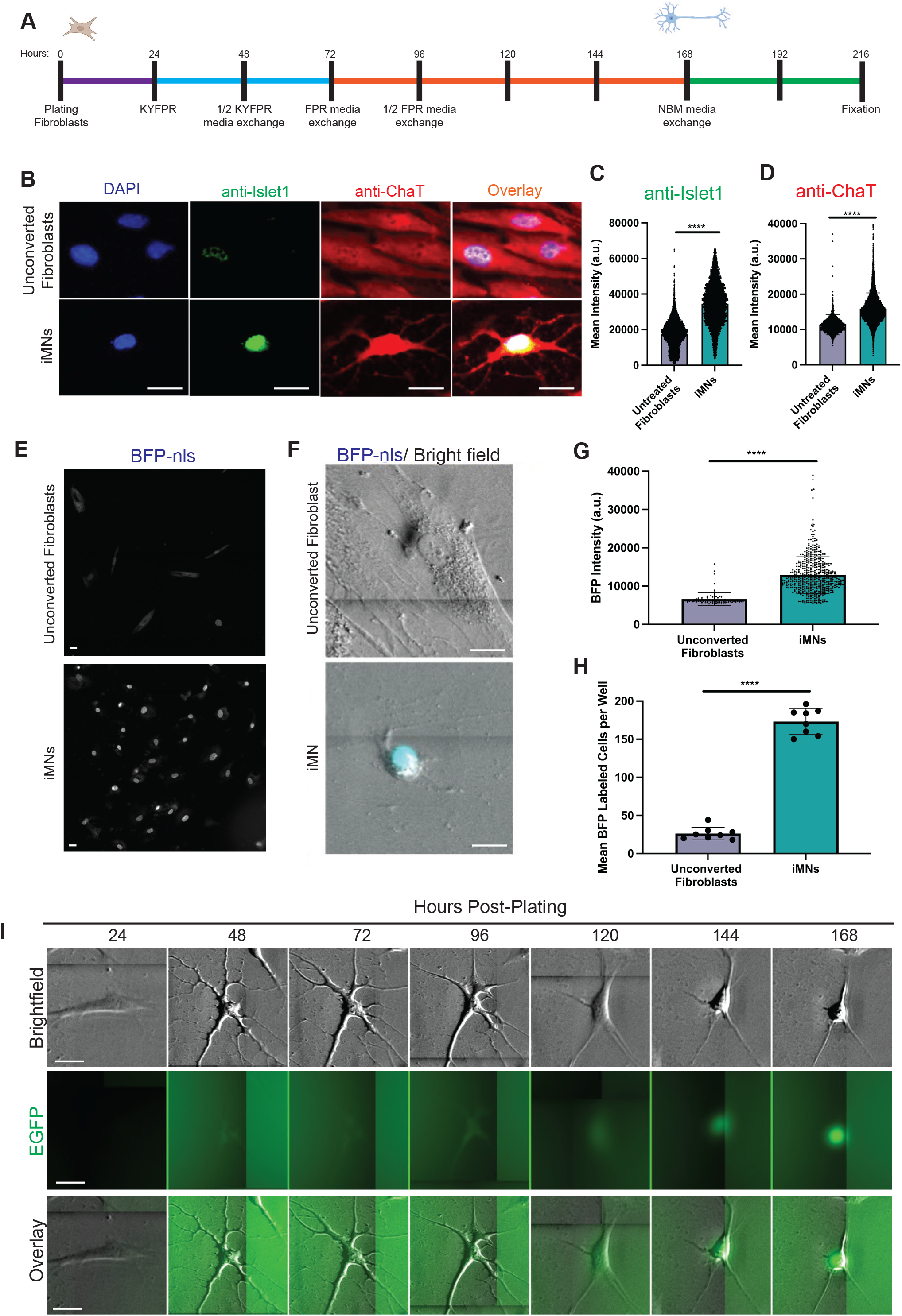
A high throughput chemical reprogramming platform for rapid genera-tion of iMNs. A) Schematic diagram of the chemical cocktail reprogramming protocol (K, kenpaullone; F, forskolin; Y, Y-27632; P, purmorphamine; R, retinoic acid). All steps were performed with automated liquid handling. B) Converted iMNs express motor neuron markers islet1 and ChAT. Scale = 25 μm. Quantification of anti-islet1 signal (C) and anti-ChAT (D) signal in iMNs compared to unconverted fibroblasts (T test ****, p<0.0001). E) Expression of BFP in iMNs and unconverted fibroblasts transduced with hSyn1:BFP-NLS. Scale = 25 μm. F) BF and BFP channel overlay of a single unconverted fibroblast and iMN transduced with hSyn1:BFP-NLS. Scale = 25 μm. Quantification of BFP intensity (G) and BFP labeled cells per well (H) in unconverted fibroblasts and iMNs transduced with hSyn1:BFP-NLS (T test ****, p<0.0001). Each replicate wells was plated with 500 fibroblasts per well. I) Morphological changes in BF and expression of EGFP during reprogramming of neuron transduced with hSyn1:EGFP. Scale = 25 μm.

### Reprogrammed iMNs retain an age signature from fibroblast donors

Although reprogramming technologies have enabled the ability to study neurological and neurodegenerative diseases from patient-specific cells, induction of pluripotency in somatic cells is thought to erase the age signature of donor cells to an embryonic state^7,26^. This greatly limits the ability to study age-related neuronal diseases such as neurodegenerative disease, where hallmarks of the disease may not manifest until later in life. Often, iPSC-derived neurons from patients with disease do not show the same neurodegenerative hallmarks (such as neuronal death) in culture without the application of stressors to the cells^4,5^. One potential reason for the lack of disease signature in iPSC-derived neurons is that they appear to have an embryonic age signature regardless of the age of the original donor^27,28^. Direct reprogramming of fibroblasts to neurons is thought to retain age signature^7^, and studies on a range of neurodegenerative diseases modeled using directly reprogrammed cells demonstrated hallmarks of disease in cellular models that have not been evident from conventionally reprogrammed iPSC-derived cells^8,29,30^. To date, retention of age signature of the fibroblast donor has only been demonstrated with viral direct reprogramming protocols^6,7^.

Methylation signature is a well-established methodology for predicting the biological age of cells^34^. Using a DNAge methylation signature clock (Zymo), we found a sequential increase in predicted age of fresh mouse primary fibroblasts, unconverted primary fibroblasts, and chemically reprogrammed iMNs that corresponded to the increased chronological age of the donor mice (Figure 2A). This data suggests a similar age-related methylation signature after culturing and reprogramming fibroblasts, and that age signature was not lost during the reprogramming protocol.

**Figure 2.**
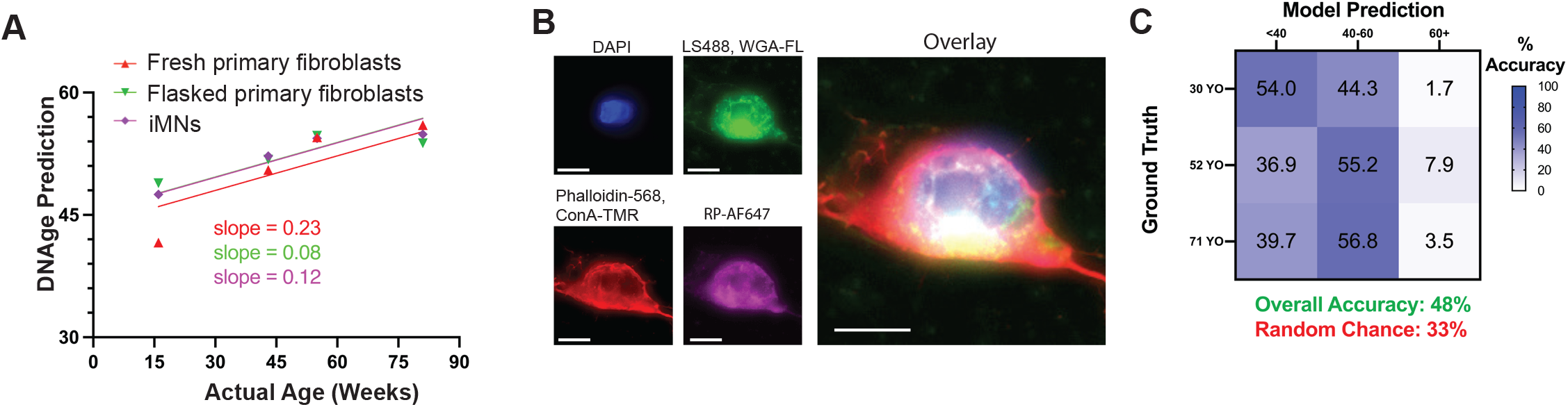
iMNs retain age signature of fibroblast donor. A) DNAge predictions from fresh mouse primary dermal fibroblasts, mouse dermal fibroblasts cultured in a flask, and mouse fresh dermal fibroblasts converted into iMNs derived from 15-, 45-, 55-, and 80-week-old mice. Lines represent linear regressions of the data used to derive slopes. B) Representative images of medium age iMNs stained with aging staining panel of DAPI, LipidSpot488, Phalloidin CF568, Wheat Germ Agglutinin-FL, Concanavalin A-TMR, and Ribosomal Protein AF647. C) Confusion matrix of deep learning model trained to predict age class tested on cell lines/iMNs model was not trained with. Scale = 10 μm.

We next attempted to detect a phenotypic age signature in images of iMNs derived from human primary fibroblast lines. A small bank of primary fibroblast lines from 10 healthy human donors across a spectrum of ages was assembled and expanded (Table 1). Induced motor neurons derived from these healthy fibroblasts were fixed and stained with a panel of morphological stains. Microscopy of these iMNs across 4 fluorescence channels produced a large, DL-compatible dataset (Figure 2B). Cells were segmented using the DAPI stain with a custom-trained, neural-network-based segmentation model. Each image was cropped into single-cell images that were used to train a convolutional neural network with the task of classifying of an iMN as young, middle, or old (see Materials and Methods^31^). The model achieved a classification accuracy of 84% on held out data randomly sampled from across experimental plates. To further validate the model, we deployed it on iMNs of a variety of ages derived from fibroblast lines that the model had not previously seen. We observed an above-chance accuracy of the model on this new dataset (48%, Figure 2C), indicating our model can detect phenotypic age signature of iMNs from donor fibroblast lines on which the model had not been trained. These data demonstrate that chemically reprogrammed iMNs retain age signature of the fibroblast donor and suggest that iMNs can model neuronal aging.

**Table 1.**
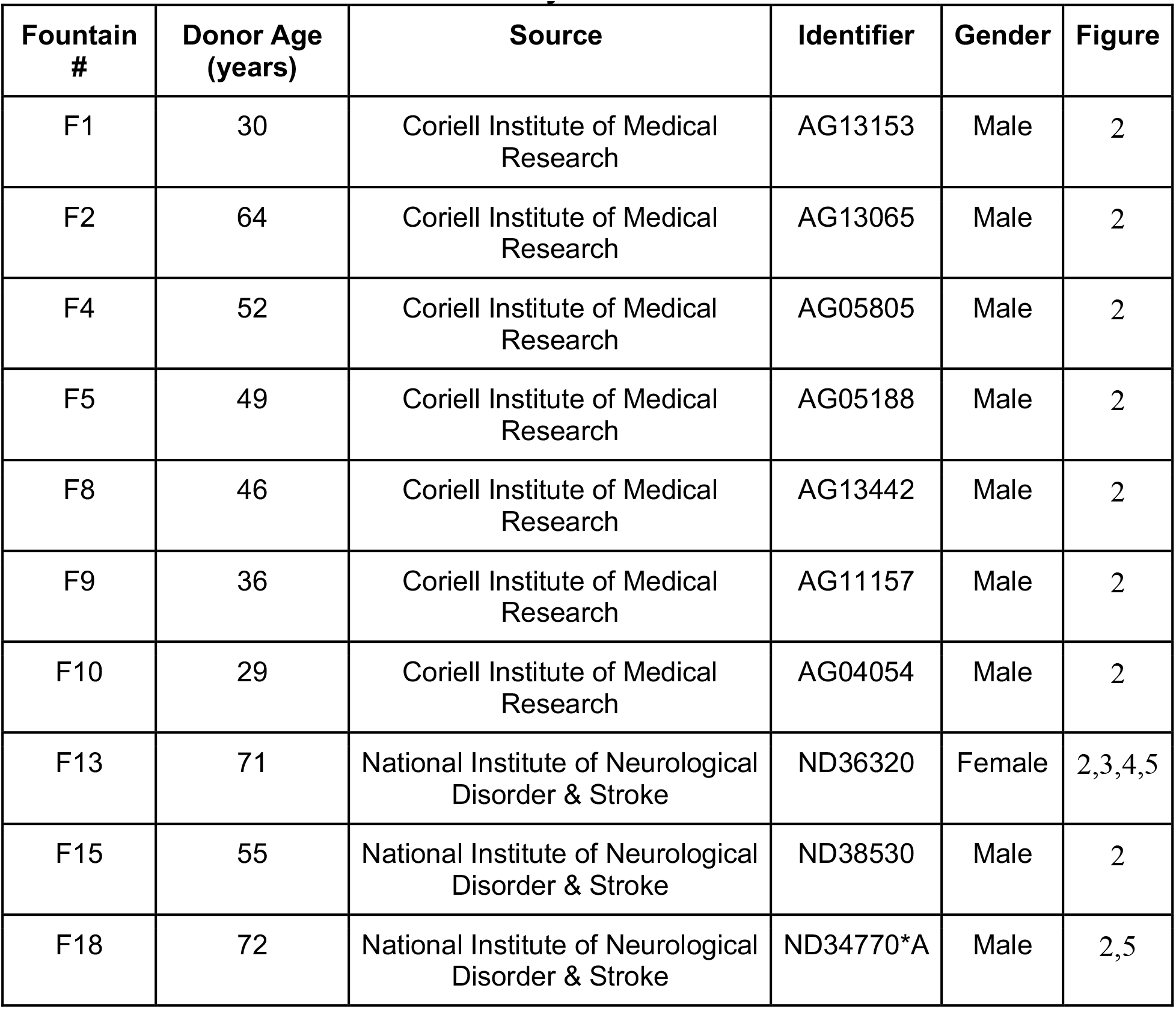
Fibroblast Lines Used in Study.

| **Fountain #** | **Donor Age (years)** | **Source** | **Identifier** | **Gender** | **Figure** |
| --- | --- | --- | --- | --- | --- |
| F1 | 30 | Coriell Institute of Medical Research | AG13153 | Male | 2 |
| F2 | 64 | Coriell Institute of Medical Research | AG13065 | Male | 2 |
| F4 | 52 | Coriell Institute of Medical Research | AG05805 | Male | 2 |
| F5 | 49 | Coriell Institute of Medical Research | AG05188 | Male | 2 |
| F8 | 46 | Coriell Institute of Medical Research | AG13442 | Male | 2 |
| F9 | 36 | Coriell Institute of Medical Research | AG11157 | Male | 2 |
| F10 | 29 | Coriell Institute of Medical Research | AG04054 | Male | 2 |
| F13 | 71 | National Institute of Neurological Disorder & Stroke | ND36320 | Female | 2,3,4,5 |
| F15 | 55 | National Institute of Neurological Disorder & Stroke | ND38530 | Male | 2 |
| F18 | 72 | National Institute of Neurological Disorder & Stroke | ND34770*A | Male | 2,5 |

### An age-relevant ALS model of iMNs shows hallmarks of disease

To model ALS, we designed lentiviral-based, neuron-specific expression constructs to transduce iMNs with blue fluorescent protein (BFP) fused to an autosomal dominant TDP mutation (TDP43^M337V^), a well-established driver of strong ALS-like phenotypes such as increased neuronal death^21,22^ (Figure 3A). Overexpression of TDP43 can also result in ALS-like phenotypes in cells and animal models^22,32^. For a control, we transduced cells with lentivirus expressing a nuclear localization sequence (NLS) fused to BFP to match the nuclear localization of TDP43 in healthy cells and avoid pathologies associated with WT TDP43 overexpression (Figure 3A). Enhanced green fluorescence protein (EGFP) was used to label cell morphology in both TDP43^M337V^-BFP-and BFP-NLS-transduced iMNs, and tracked with longitudinal imaging every 6 hours in neural basal media containing B27plus (Thermo), but without additional growth factors. Neuronal death could be observed by the loss or reduction of BFP or EGFP fluorescence as previously described^33^, but was also particularly obvious in the brightfield channel (Figure 3B), where dead cells showed a clear loss of membrane birefringence and clumping in the soma. Cells were tracked over time and scored manually for death. Survival statistics were fit with a Kaplan-Meier curve, and a Cox proportional hazards model was derived. Results show that iMNs expressing TDP43^M337V^-BFP had a higher likelihood of death than control neurons expressing BFP-NLS, indicating TDP43^M337V^ overexpression in iMNs may induce ALS-related phenotypes (Figure 3C).

**Figure 3.**
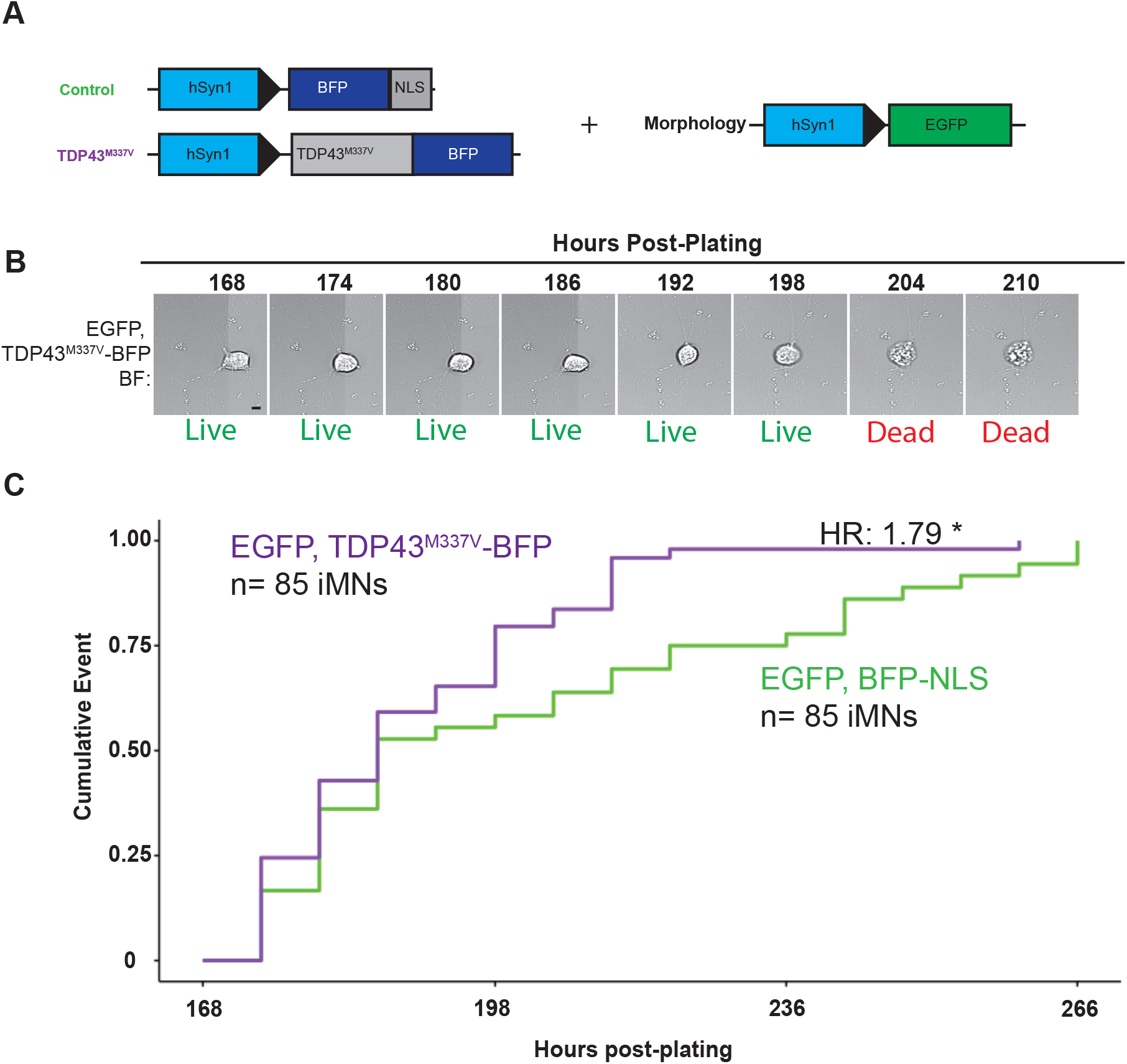
Transduction of TDP43-BFP reduces survival of iMNs. A) Schematic figure of control and TDP43^M337V^-BFP constructs co-transduced with hSyn1:EGFP used to model TDP43-ALS. B) Time lapse BF imaging of degenerating iMN co-transduced with hSyn1:TDP43^M337V^-BFP and hSyn1:EGFP. C) Kaplein-Meyer survival plots of neurons co-transduced with hSyn1:TDP43^M337V^-BFP and hSyn1:EGFP Cox proportional hazard 1.79, *,p<0.05. Scale = 10 μm.

Next, a screening platform was set up to identify small molecules that revert TDP43^M337V^-induced pathology. Previous screening protocols have focused on the degree to which TDP43 mutation leads to loss of TDP43 nuclear localization and an increase in TDP43-positive cytoplasmic aggregates, features that correlate with known disease pathology^22,34^. Yet gross quantification of non-nuclear localization of TDP43 likely over-simplifies the complexity of TDP43-associated pathology^35^. For instance, quantification of TDP43 localization fails to account for the onset of positive compensatory responses to pathological TDP43 that are likely to occur outside the nucleus^22^, the pathological functions of TDP43 that occur within the nucleus^36,37^, or the localization of TDP43 aggregation^38,39^. Deep learning image analysis techniques offer the ability to decode such complex spatial-phenotypic patterns in a more sophisticated way^40^.

We next attempted to train DL models capable of detecting ALS-associated pathology that we could probe with small molecules that might revert the phenotype back to a normal state. Large training datasets of iMNs generated from a fibroblast cell line derived from a 71-year-old healthy donor were transduced with either TDP43^M337V^-BFP or BFP-NLS. Individual cells were segmented based on their BFP signal using a custom nuclear segmentation algorithm. Crops of individual cells were generated and filtered out if they contained BFP intensity below an arbitrary intensity threshold to eliminate un-converted fibroblasts (Figure 1E–H). BFP channel single cell images were used to train a model to classify whether a cropped image of a cell is either TDP43^M337V^-BFP or BFP-NLS control (Figure 4A). Testing on randomly held out images showed high accuracy per image (Figure 4B–C). Nevertheless, BFP expression within iMNs could be affected by experimental artifacts such as differences in viral expression levels unrelated to pathology induced by the overexpressed TDP43^M337V^ mutation. To test if differences in TDP43^M337V^-BFP and BFP-NLS control transduced cells could be observed outside of BFP expression levels, we stained iMNs in the training set with WGA-647, a disease-agnostic membrane stain, and reimaged iMNs with far red (WGA-647) and transmitted light/brightfield (BF) channels (Figure 4D). A new deep learning model trained on WGA- 647 and BF channels showed similar accuracy, indicating broad phenotypic differences between TDP43^M337V^-BFP and BFP-NLS control transduced cells (Figure 4E). These results indicate TDP43^M337V^-BFP viral transduced iMNs manifest phenotypes capable of being detected by deep learning models trained against TDP43 signal as well as unbiased, disease-agnostic labels such as WGA-647 and BF.

**Figure 4.**
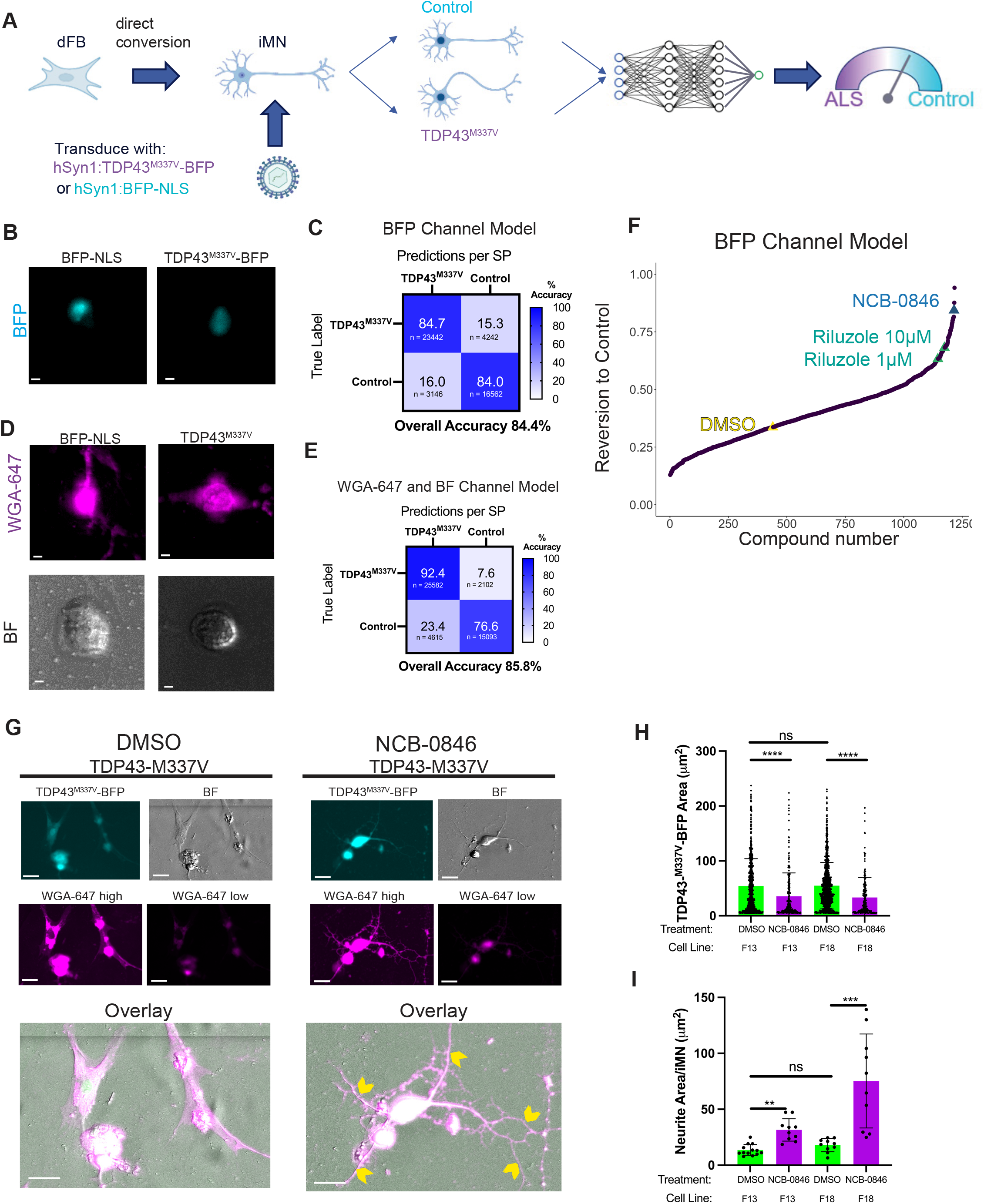
Small molecule screening of TDP43-ALS iMNs leads to identification of NCB-0846. A) Schematic figure of development of a DL model for identifying TDP43- ALS phenotypic signatures. Converted iMNs are transduced with hSyn1:TDP43^M337V^- BFP or hSyn1:BFP-NLS control, imaged, and used to train a DL model capable of identifying compounds that revert ALS phenotypes back to control phenotypes. B) Representative images of BFP-NLS or TDP43^M337V^-BFP transduced iMNs. Scale = 10μm. C) Confusion matrix of performance of DL model trained to predict TDP43^M337V^ or control using BFP channel. D) Representative images of TDP43^M337V^-BFP or BFP-NLS control transduced neurons stained with WGA-647 and imaged in brightfield or far red channel. Scale = 10 μm. E) Confusion matrix of performance of DL model trained to predict ALS or control using BF and WGA-647 channels. F) Scoring of compound library for reversion to control by BFP channel ALS vs control model. G) Representative images of DMSO treated and NCB-0846 treated iMNs stained with WGA-647. High and low contrast 647 channel images are displayed to highlight neurite detail and high dynamic range of WGA-647. Yellow chevrons indicate increased neurite branching. Scale = 25 μm. H) Quantification of TDP43^M337V^-BFP area in iMNs generated from two different cell lines and treated with DMSO or NCB-0846. ANOVA, Tukey’s **** p<0.0001, ns = not significant. I) Quantification of neurite area per iMN of TDP43^M337V^-BFP area in iMNs generated from two different cell lines and treated with DMSO or NCB-0846. ANOVA, Tukey’s *** p<0.001, ** p<0.01, ns = not significant.

Next, we tested whether molecules that reversed aging in dermal fibroblasts (dFBs) (data not shown) showed a therapeutic effect in our TDP43^M337V^-BFP vs BFP-NLS neuron assay. Converted iMNs expressing TDP43^M337V^-BFP were treated with a library of 1009 small molecules that previously showed a reduction in age in an internal screen in dFBs (data not shown) in replicate at 10 μM and imaged in the BFP channel. Each imaged iMN was given a predicted class score ranging from 0 as TDP43^M337V^ to 1 as control. The average score across all iMNs within a well was calculated, and then averaged across well replicates. A positive control, riluzole, one of the few FDA-approved for treatments for ALS that extends the lifespan of patients^44^ but which has an elusive mechanism of action^41,42^, showed a strong efficacy at both 1 (randomization test p<0.0001) and 10μM (randomization test p<0.0001) concentrations in reverting the phenotype of TDP43^M337V^-BFP towards control (Figure 4F). DMSO, the negative control, showed no efficacy at reverting the phenotype back to control. Three compounds showed a reversion of 3 standard deviations above the mean, one of which was NCB- 0846 (randomization test p<0.0001), a TNIK/MAP4K7 inhibitor^43^. Induced motor neurons expressing TDP43^M337V^-BFP treated with NCB-0846 showed clear, morphological changes including a more compact nuclear TDP43^M337V^-BFP signal and increased neurite branching as seen when labeled with WGA-647 (Figure 4G). To validate these changes, fibroblasts from a different normal (non-ALS) human donor of similar age (72 years old, Table 1, cell line “F18”) were converted to iMNs in parallel and showed similar significant decreases in TDP43^M337V^-BFP area and increases in neurite branching, suggesting increased neuronal health and decreased non-nuclear TDP43 expression (Figure 4H–I). These data support further investigation of NCB-0846 as a potential candidate molecule for reverting ALS-like phenotypes in motor neurons.

### NCB-0846 shows therapeutic efficacy against a SOD1^G93A^ age-relevant iMN model

We next developed the assay using a model of SOD1 driven ALS. Mutations in SOD1 are the most common familial mutations that drive ALS, though they are thought to result in unique and possibly non-overlapping pathogenetic processes and may not result in TDP43 pathology that occurs in over 90% of ALS cases^35^. Dominant negative mutations in SOD1 such as G93A are the second most common cause of familial ALS, and overexpression of SOD1^G93A^ in cellular neuronal models has been previously de-scribed to result in ALS-like hallmarks such as increased neuronal death^23^. We designed lentiviral-based, neuron-specific expression constructs to transduce iMNs with SOD1^G93A^ fused via a P2A cleavable peptide to EGFP, or EGFP alone as a control, and generated a large dataset of iMNs stained with DAPI and WGA-647 (Figure 5A). Imaging was performed using DAPI, EGFP, WGA-647 and brightfield channels. Induced motor neurons were segmented using the nuclear DAPI signal. As previously, single-cell images of iMNs with low EGFP signal were filtered out to reduce unconverted fibroblast contamination. Single-cell, four-channel images of iMNs were used to train a DL model to classify iMNs as either SOD1^G93A^-P2a-EGFP or EGFP control (Figure 5B–D). Trained models showed high accuracy (Figure 5E). Next, we screened the same 1009 molecules previously screened for TDP43^M337V^-BFP versus BFP-NLS control. NCB-0846 again scored as a hit in the screen (>2 SD above the mean, randomization test p<0.0001) (Figure 5F). Induced motor neurons treated with NCB-0846 reverted SOD1^G93A^ phenotypes to control better than the positive control riluzole (randomization test p<0.0001) and displayed increased neurite branching (Figure 5F–G). Additionally, another TNIK/MAP4K7 inhibitor, KY-05009^44^, was added to the screen and scored even higher than NCB-0846 (randomization test p<0.0001) (Figure 5H). These data reinforce the therapeutic potential of TNIK/MAP4K7 as a therapeutic target for ALS.

**Figure 5.**
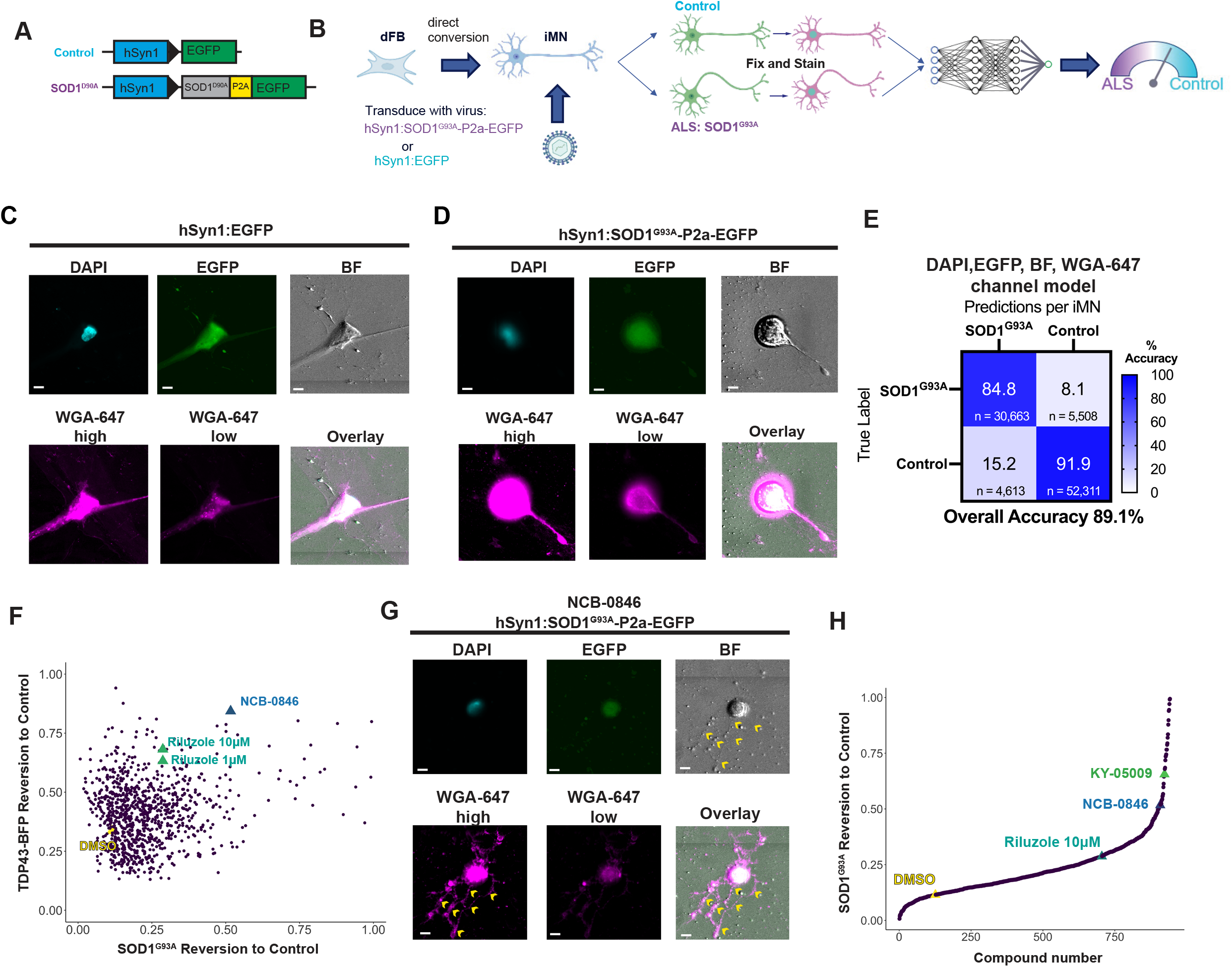
Small molecule screening of SOD1-ALS iMNs confirms NCB-0846 effi-cacy towards ALS. A) Schematic figure of control and SOD1^G93A^ constructs used to model SOD1-ALS. B) Schematic figure of the development of a deep learning model for identifying SOD1-ALS phenotypic signatures. Converted iMNs are transduced with hSyn1:SOD1^G93A^-P2a-EGFP or hSyn1:EGFP control, fixed and stained with WGA-647 and DAPI, and images are used to train a deep learning model capable of identifying compounds that revert SOD1-ALS phenotypes back to control phenotypes. C) Repre-sentative images of iMNs transduced with hSyn1:EGFP or D) hSyn1:SOD1^G93A^-P2a-EGFP, stained with DAPI and WGA-647, and imaged in DAPI, EGFP, far red, and BF channels. E) Confusion matrix of performance of deep learning model trained to predict SOD1-ALS or control using DAPI, EGFP, far red, and BF channels. F) Correlation of scoring between TDP43-ALS reversion to control and SOD1-ALS revision to control models on chemical library highlighting DMSO (negative control), riluzoler (positive con-trol), and NCB-0846. G) Representative images of iMNs transduced with hSyn1:SOD1^G93A^-P2a-EGFP treated with NCB-0846, stained with DAPI and WGA-647, and imaged in DAPI, EGFP, far red, and BF channels. High and low contrast 647 chan-nel images are displayed to highlight neurite detail and high dynamic range of WGA- 647. Yellow chevrons indicate increased neurite branching. H) Scoring of SOD1-ALS compound reversion to control for compound library highlighting DMSO, Riluzole, NCB- 0846, and TNIK/MAP4K7 inhibitor KY-05009. Scale = 10 μm.

### NCB-0846 shows therapeutic potential against an ALS mouse model

We next sought to test the efficacy of NCB-0846 in a commonly used ALS mouse model, SOD1^G93A^ ^45^. Hemizygous female mice expressing the SOD1^G93A^ transgene were administered NCB-0846 or vehicle control via oral gavage at a single dose of 30 mg/kg beginning at 10 weeks of age. Animals dosed with NCB-0846 did not show improvements in wire hang or rotarod assays during treatment (Figure 6A–B), though the experiment was likely underpowered with only 6 mice per condition and a single dose concentration. At the terminal endpoint, serum was collected from mice and assayed for levels of Neurofilament Light Chain (NfL), an intracellular protein and biomarker for axonal damage that is released into the cerebral spinal fluid and blood on cell death (citation). NfL levels are known to increase in both patients and mouse models of ALS^46^, and NCB-0846 treated mice showed significant reduction in levels versus vehicle treated SOD1^G93A^ mice (Figure 6C).

**Figure 6.**
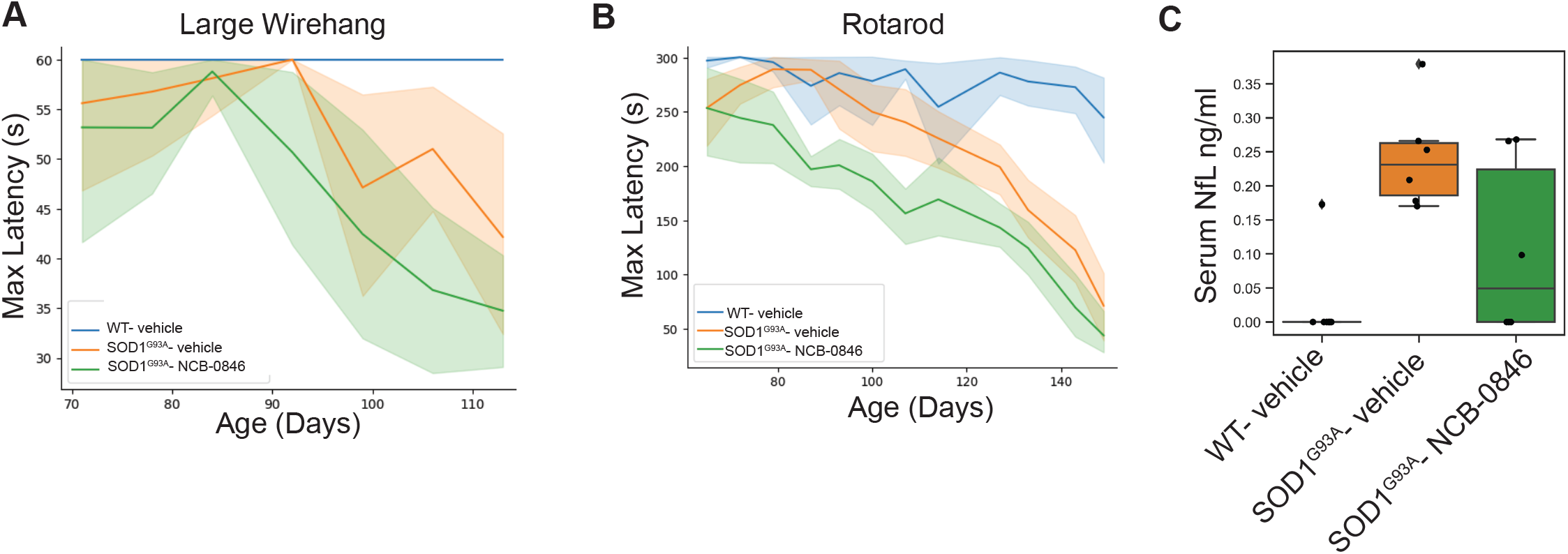
NCB-0846 administration reduces NfL levels in SOD1^D93A^ mice. A) NCB- 0846 administration does not improve large wire hang, or B) rotarod performance. C) NfL levels are significantly reduced in mice administered NCB-0846. One-Way ANOVA with Multiple Comparisons (Tukey’s correction) *** p<0.001, * p<0.05.

## Discussion

The complex and age-related nature of neurodegenerative diseases like ALS have long proved challenging for study, contributing to the paucity of treatments for these diseases. To start, animal models often fail to recapitulate key aspects of age-related neurodegenerative diseases and therapeutic improvements in animals have not translated well to the clinic^47^. The recent advent of cellular reprogramming and directed differentiation technologies have enabled the generation of neurons from patient-specific iPSCs. These iPSC-derived neurons hold great promise for modelling and studying neurological development and diseases; however, neurons thus generated are age-re-set to an embryonic stage^6,7,29,48,49^, and do not recapitulate the aging component of neurodegenerative diseases. Finally, the process of aging itself introduces an immense amount of complexity to biology, as different cells and tissues within the same organism often age at different rates^50^. This effect introduces increased variability, which complicates many aging studies^55^. Thus, challenges generating models of neurodegenerative diseases with their associated phenotypes have limited our ability to find much-needed treatments for these diseases. To address some of these challenges, we created an screening platform to generate and take advantage of image-based machine learning techniques. We then used this platform to analyze age-relevant human motor neurons with ALS-like pathology to identify small molecules with therapeutic efficacy in both *in vitro* and *in vivo* ALS models.

Notably, our screens identified the TNIK/MAP4K7 inhibitor NCB-0846 as a molecule with the ability to revert TDP43 and SOD1 mutant phenotypes *in vitro* and with potential disease modifying effects *in vivo*. NCB-0846 has been previously shown to have efficacy in other indications that involve the TNIK/MAP4K7 pathway, including colorectal neoplasms^43^, lung cancer^51^, synovial sarcoma^52^, and hepatic fibrogenesis^58^, but not ALS. Inhibitors of another MAPK4 protein, MAP4K4, have previously been shown to exhibit efficacy in ALS models, and MAP4K inhibitors have also been evaluated as neuron protecting agents in iPSC derived motor neurons^53–56^. MAP4Ks belong to the GCK-IV family of the Ste20 group kinases^57,58^, and share similar protein structures with a high homology within the kinase domain^59^. Thus, many small molecules that inhibit one may also inhibit other MAP4K family members, and it remains possible that the effects we observe from NCB-0846 and KY-05009 could result from inhibition of one or more MAP4Ks besides MAP4K7. A recent non-peer-reviewed report^66^ described MAP4Ks as a therapeutic target for ALS by using a viral-based direct reprogramming strategy to generate iMNs, suggesting MAP4Ks may have more relevance in ALS motor neuron models where age signature is retained. Interestingly, that report found that inhibition of specific MAP4Ks, in particular MAP4K4, MAP4K6, and MAP4K7, can prolong survival of iMNs, and may activate a novel downstream signaling pathway in aging-relevant neurons not found in studies using embryonic iPSC-derived neurons. Therefore, our discovery of MAP4K7 inhibitors NCB-0846 and KY-05009 as having efficacy in ALS models, could reflect key biological differences between embryonic and age-relevant neurons.

One major advance in our work was the use of chemical reprogramming for HTS. Here, we took advantage of the rapid time course of reprogramming, age-retention, and reprogramming efficiency of the motor neuron chemical reprogramming protocol to conduct an HTS that is compatible with DL image analysis techniques. Chemical reprogramming methodologies now exist for converting fibroblasts into other cell types such as dopaminergic neurons^60^, astrocytes^61^, monocytes^69^, cardiac myocytes^62^, macrophages^71^, and microglia^63^, suggesting this HTS screening platform could be adopted towards a plethora of tissue-specific indications other than just motor neurons/ALS to generate age-relevant cells and identify therapeutic compounds. Additionally, our ability to identify an age signature in motor neurons suggests we may be able to discover small molecules that generally reverse aging in neurons that may have positive impact across multiple specific disease contexts.

In summary, we developed a deep-learning-powered screening platform for identifying small molecule therapeutics in age-relevant human ALS models. Our screen identified a neuroprotective chemical compound, NCB-0846, which when tested further was shown to have potential disease-modifying effects in additional *in vitro* and *in vivo* ALS models. Future studies to expand in vivo testing of NCB-0846 at different doses and with larger mouse cohorts are needed, as is testing of more selective MAPK4 family inhibitors, to better identify potential disease modifying targets and effects in motor neurons. Screening of larger and more diverse chemical libraries in this screening platform should yield additional hits that may reverse the effects induced by the expression of ALS-associated mutant proteins, and/or reduce age-relevant effects that allow diseases such as ALS to be more penetrant in aged motor neurons.

## Materials and Methods

### Human primary Fibroblast Culture and iMN reprogramming

Human fibroblast lines were acquired from Coriell, and expanded in fibroblast growth media (Dulbecco’s modified eagle medium-DMEM, 15% fetal bovine serum, 1:100 non-essential amino acids, 1:100 sodium pyruvate, 1:100 Glutamax, 1:100 Hepes, 1:100 Beta-Mercaptoethanol, 1:100 Pen/Strep) before being banked in liquid nitrogen. After thawing, fibroblasts were allowed to grow at least two weeks in culture in fibroblast growth media in plastic flasks before assaying.

Glass bottom 384-well plates were coated with poly-D-lysine and matrigel diluted 1:100ml in chilled DMEM/F-12 and incubated at 37degrees at least overnight. When 90% confluent, fibroblasts were trypsinized and plated onto coated plates at 1000 cells per 384 plate well using an Integra Viaflow384. Fibroblasts were allowed to settle overnight. iMNs were then converted from adult human skin fibroblasts as previously de-scribed with modifications^25^. Briefly, the day after seeding, the fibroblast media was exchanged after a wash with PBS into Kenpaullone (20 μMM), Y27632 (20 μMM), Forskolin (20 μMM), Purmorphamine (20 μMM), Retinoic acid (40 μMM) cocktail in neurobasal media containing media containing normacin, glutamax, and B27plus. A 50% media change was performed the following day. The media was then exchanged for Forskolin, Purmorphamin, Retinoic Acid (FPR) in neurobasal media containing media containing normacin, glutamax, and B27 plus the following day. The next day, viral transduction was performed in FPR media and incubated three more days. Next, media was exchanged into screening media (NBM, normacin, glutamax, and B27 plus). All media exchanges were performed using a BioTek plate washer. Converted iMNs were then fixed and stained after two day incubation in screening media. Cells were fixed with 10% Neutral Buffered Formalin (Leica 3800600) for 20 minutes and washed with PBS with 50 μgg/mL gentamicin. For survival assay, screening media was changed daily.

### Plasmids and virus production and drug treatment

Coding sequence for hTDP43^M337V^-BFP, BFP-NLS, hSOD1^G93A^, and EGFP were synthesized by GenScript, and subcloned into pGenLenti backbone. Lentivirus was generated by GenScript or BrainVTA with titer QC’d at >1×10e8TU10e8TU/mL. Virus was applied to wells with 1:100 dilution.

### Imaging, image processing, and model training

All imaging was performed on Celldiscoverer 7 (Zeiss) high content imager with 20X/0.95 NA objective, Colibri 7 LEDs, and Axiocam 506 camera and Zen 3.1 software. Robotic arms were used to load individual plates from a stack into the CD7 using integrated plate loading software (Let’s go robotics). Images were collected with 5×5 tiling per well and were saved in .czi format per plate imaged. The outer edges of 384 well plates were not imaged due to well established plate effects, and only 264 wells are used per plate. Images were processed into czi’s per individual well, and then segmented using a custom trained U-net segmentation algorithm trained with custom annotations on the DAPI stains. Individual 202×202 crops were generated for each cell and uploaded to the S3 cloud for storage and processing.

### Antibody staining and cell signal quantifications

For age-prediction experiments, staining was performed in two steps. First, an overnight incubation at 4C, with 0.1% Triton X-100, anti-SC35 Alexa Fluor 488 (1:6000), and anti-S6 Ribosomal Protein Alexa Fluor 647 (1:2000). Next, a room temperature, 45 minutes incubation with DAPI (1:1000), LipidSpot 488 (1:3000), wheat germ agglutinin fluorescein (1:2500), phalloidin CF568 (1:166.67), and Concanavalin A Tetramethylrhodamine (1:833.33). DAPI was prepared at 1 μgg/mL in a 1 L glass bottle. Fixed and DAPI stained plates were stored at 4°C prior to imaging for up to five days.

iMNs were stained with rabbit anti-ISL1 (1:200, ab20670, Abcam), goat anti-ChAT (1:200, AB144P, Millipore), Alexa 488- and Alexa 594- conjugated secondary antibodies (LifeTechnologies), and DAPI. Cell segmentation was performed on DAPI signal using intensity and size thresholds and derived masks were used to quantify the mean intensity signal across channels. For lentiviral quantification, TDP43^M337V^-BFP or BFP-NLS signal was used for segmentation and derived masks were used for quantification of area, intensity, or labeled cell count. Neurite quantification was performed using NeurphologyJ on randomly sampled 1500×1500 crops of the WGA-647 signal.

### Model training

For each of the models -- iMN age signature model, iMN TDP43 models, and iMN SOD1 model-- we trained a modified ImageNet-pretrained ResNet-18 architecture on a single NVIDIA 3080 node for 100 epochs over training and validation sets. Training data augmentations consisted of random horizontal and vertical flips applied to each batch. For the iMN age model, we utilized 25,429 5×5 crops for training and 2,826 crops for validation from across donors F1, F2, F4, F5, F8, F9, F10 and F13. For the iMN TDP43 models, we utilized 426,525 single cell crops for training and 47,392 single cell crops for validation, all from donor F13. For the iMN SOD1 model, we utilized 837,855 single cell crops for training and 93,095 single cell crops for validation, all from donor F13.

### Statistical Analyses

For small molecule screens, randomization tests^64^ were used to estimate the probability that predicted ages for iMNs (softmax output) treated with a compound were different than control (DMSO treated). Briefly, in this procedure we first computed the difference between the mean predictions of a compound treatment versus DMSO treatment (the “true score”). Second, we simulated a null distribution of 10,000 difference scores by randomizing the association between treatment labels and their effect, then recomputing the difference between compound treatment versus DMSO treatment (“simulated scores”). Finally, we computed a p-value as the proportion of simulated scores that were greater than the true score.

Survival analysis was performed by defining the time of death as the point at which a neuron appeared to have a clear loss of membrane birefringence and clumping in the soma. The survival package for R statistical software was used to construct Kaplan–Meier curves from the survival data, and survival functions were fit to these curves to derive cumulative survival and risk-of-death curves that describe the instantaneous risk of death for individual neurons as previously described^65^.

### Mouse Experiments

SOD1^G93A^ transgenic mice (Stock #004435, B6.Cg-Tg(SOD1*G93A)1Gur/J) and C57BL/6J (Stock #000664) mice were obtained from The Jackson Laboratory (Bar Harbor, ME). All mice were housed in temperature-and humidity-controlled rooms (22 ± 2 °C and 35–65%, respectively) with a 12-hour light/12-hour dark cycle, under specific pathogen free conditions in CRADL (Charles River Accelerator and Development Lab, 750 Gateway Blvd., South San Francisco, CA). All animal procedures were followed in compliance with the Institutional Animal Care and Use Committee (IACUC) at Charles River Laboratories.

Beginning at 10 weeks of age, mice were dosed with Vehicle or NCB-0846 (Med-Chem Express, Catalog # HY-100830) via p.o. 5-daily at 30 mg/kg in (PEG400:Tween-80:water). Mice were dosed until 22 weeks of age or until humane endpoint. Mice were anesthetized with 2.5% Avertin^73^ (2,2,2-tribromoethanol, Sigma T48402, solubilized in 2-Methyl-2-butanol, Sigma 152463, and dissolved to a working solution 1:40 with 1X PBS) via i.p. injection at 250 mg/kg body weight. Blood was collected via intracardial puncture with a 26G x 1/2 needle (BD 305111), placed into Serum Separator tubes (Greiner Bio-One, MiniCollect Serum Tubes, #450472) and kept on ice. After blood collection, mice were euthanized via cervical dislocation. Lumbar spine, legs, brain, and eyes were collected and fixed in 4% PFA. Blood samples were centrifuged at 1000 G for 15 min at 4 °C and stored at -80 °C. Neurofilament light chain (NF-L) was quantified using Mouse NF-L ELISA Kit (Novus Biologicals, Catalog #NBP2-80299) and read on a SpectraMax iD5 (Molecular Devices, San Jose, CA).

For DNAge experiments, primary dermal fibroblasts from 3-, 9-, 12-, and 18- month-old C57BL/6J (Stock #000664) mice were isolated. Animals were CO_2_ euthanized and dissected skin samples were transported in DMEM to the laboratory on wet ice. Skin samples were minced to 1-2 mm pieces and digested with Digestion Media: Collagenase (Vitacyte, Catalog #s 011-1010, 011-1060), dispase (gibco, Catalog # 17105-045), and DNase in HBSS, for 3 hours at 37 °C. A crude live cell mixture was obtained by filtration and negative selection using Easy Eights EasySep Magnet and Dead Cell Removal kit (Stemcell Technologies, Catalog # 17899). Cells were further enriched with positive CD90.2 selection (Stemcell Technologies, Catalog # 18951), washed, filtered, and resuspended in plating media. Cells were plated into collagen and poly-d-lysine coated 384-well plates, plastic 12-well culture plates, or plastic flasks for three days and then trypsinized and replated in 12-well culture plate and incubated overnight at 37 °C, 5% CO_2_, and 100% humidity. After one day recovery in plastic culture plate, iMN reprogramming was initiated, and then cells were harvested in DNAge cell lysis buffer (Zymo Research, Catalog # R1100-50), frozen, and then shipped to ZymoGenetics for CpG DNA methylation sequencing and age predictions.

### Behavior assays

At 9 weeks of age, mice were tested for wire hang (small and large) and rotarod. Mice were separated into equal groups based on their wirehang performance, rotarod performance, and weight. From 10 to 16 weeks of age, mice were tested weekly for wirehang performance. From 10 weeks of age until end of study, mice were tested weekly for rotarod performance. For the wire hang test we used a small (1mm aperture, woven 20 mesh) and a large (10 mm aperture) steel grid, where the small grid is more difficult than the large. Mice were gently placed onto the grid and inverted such that the mice were hanging upside down over a home cage. Using a manual stopwatch, the latency to fall was recorded, with a max latency of 60 seconds. Mice were tested up to 3 times per session, with a 15-minute inter-trial-interval.

In order to track changes in motor coordination, we tested animals in the rotarod test (Maze Engineers). Mice were placed onto an accelerating rotating rod and measured for ability to stay on the rod. Latency to fall, distance traveled, and speed at fall were recorded. Mice were tested for 3 trials per session, with an inter-trial-interval of 15 minutes. At 8 weeks of age, mice were habituated to the rotarod at a constant 4 RPM for a max latency of 5 minutes. Beginning at 9 weeks of age, mice were tested on an acceleration protocol. Rotation would begin at 4 RPM and accelerate at 7.2 RPM over 5 minutes to a final 40 RPM.

## Declarations

### Authors contributions

Jalnapurker B and Linsley JW cultured cells, stained cells, and performed iMN reprogramming. Pipathsouk A and Linsley JW performed high throughput microscopy. Martinez F. Curt C., Rodgers J., Corbin C., developed image processing pipelines and processed image data. Shainin W., and Zarconne R built machine learning models. Linsley JW, Bhatnagar R., Rosen J., Rodgers J., and Le D. performed data analysis. Bhatnagar R. Rosen J., and Rodgers J., generated compound library. Le D. and Rodgers J., performed animal studies. Hodge B., Martinez F., Pipathsouk A., Chin J., Jalnapurker B, Rodgers J., and Linsley JW., performed laboratory experiments. All authors discussed results / reviewed the manuscript. Linsley JW drafted the original manuscript, conceived of, and designed experiments. Linsley JW, Rodgers J., Rando TA, and Zar-cone R. oversaw the project.

## Acknowledgements

We thank William Greene, Anu Hoey, Lynn Yamamoto, Vinh Tran, and Derek Britain for helpful discussion and technical assistance. We thank Erin Daruszka for administrative assistance. We thank Drew Linsley for guidance on statistical modeling and machine learning. Illustrations were generated using BioRender.com.

## References

1 Ingre, C., Roos, P. M., Piehl, F., Kamel, F. & Fang, F. Risk factors for amyotrophic lateral sclerosis. Clin Epidemiol 7, 181–193, doi:10.2147/clep.S37505 (2015).

2 Pandya, V. A. & Patani, R. Decoding the relationship between ageing and amyotrophic lateral sclerosis: a cellular perspective. Brain 143, 1057–1072, doi:10.1093/brain/awz360 (2020).

3 Brown, R. H. & Al-Chalabi, A. Amyotrophic Lateral Sclerosis. N Engl J Med 377, 162–172, doi:10.1056/NEJMra1603471 (2017).

4 Linares, G. R. et al. SYF2 suppression mitigates neurodegeneration in models of diverse forms of ALS. Cell Stem Cell 30, 171–187.e114, doi:10.1016/j.stem.2023.01.005 (2023).

5 Workman, M. J. et al. Large-scale differentiation of iPSC-derived motor neurons from ALS and control subjects. Neuron 111, 1191–1204.e1195, doi:10.1016/j.neuron.2023.01.010 (2023).

6 Huh, C. J. et al. Maintenance of age in human neurons generated by microRNA-based neuronal conversion of fibroblasts. Elife 5, doi:10.7554/eLife.18648 (2016).

7 Mertens, J. et al. Directly Reprogrammed Human Neurons Retain Aging-Associated Transcriptomic Signatures and Reveal Age-Related Nucleocytoplasmic Defects. Cell Stem Cell 17, 705–718, doi:10.1016/j.stem.2015.09.001 (2015).

8 Monk, R., Lee, K., Jones, K. S. & Connor, B. Directly reprogrammed Huntington’s disease neural precursor cells generate striatal neurons exhibiting aggregates and impaired neuronal maturation. Stem Cells 39, 1410–1422, doi:10.1002/stem.3420 (2021).

9 Pu, J. et al. Parkin mutation decreases neurite complexity and maturation in neurons derived from human fibroblasts. Brain Res Bull 159, 9–15, doi:10.1016/j.brainresbull.2020.03.006 (2020).

10 Sun, Z. et al. Endogenous recapitulation of Alzheimer’s disease neuropathology through human 3D direct neuronal reprogramming. bioRxiv, 2023.2005.2024.542155, doi:10.1101/2023.05.24.542155 (2023).

11 Pancotti, C. et al. Deep learning methods to predict amyotrophic lateral sclerosis disease progression. Sci Rep 12, 13738, doi:10.1038/s41598-022-17805-9 (2022).

12 Lee, J. et al. Deep learning-based brain age prediction in normal aging and dementia. Nat Aging 2, 412–424, doi:10.1038/s43587-022-00219-7 (2022).

13 Eulenberg, P. et al. Reconstructing cell cycle and disease progression using deep learning. Nat Commun 8, 463, doi:10.1038/s41467-017-00623-3 (2017).

14 Christiansen, E. M. et al. In Silico Labeling: Predicting Fluorescent Labels in Unlabeled Images. Cell 173, 792–803.e719, doi:10.1016/j.cell.2018.03.040 (2018).

15 Linsley, J. W. et al. Superhuman cell death detection with biomarker-optimized neural networks. Sci Adv 7, eabf8142, doi:10.1126/sciadv.abf8142 (2021).

16 Colasante, G. et al. Rapid Conversion of Fibroblasts into Functional Forebrain GABAergic Interneurons by Direct Genetic Reprogramming. Cell Stem Cell 17, 719–734, doi:10.1016/j.stem.2015.09.002 (2015).

17 Jiang, H. et al. Cell cycle and p53 gate the direct conversion of human fibroblasts to dopaminergic neurons. Nat Commun 6, 10100, doi:10.1038/ncomms10100 (2015).

18 Blanchard, J. W. et al. Selective conversion of fibroblasts into peripheral sensory neurons. Nat Neurosci 18, 25–35, doi:10.1038/nn.3887 (2015).

19 Hu, W. et al. Direct Conversion of Normal and Alzheimer’s Disease Human Fibroblasts into Neuronal Cells by Small Molecules. Cell Stem Cell 17, 204–212, doi:10.1016/j.stem.2015.07.006 (2015).

20 Sepehrimanesh, M., Akter, M. & Ding, B. Direct conversion of adult fibroblasts into motor neurons. STAR Protoc 2, 100917, doi:10.1016/j.xpro.2021.100917 (2021).

21 Barmada, S. J. et al. Autophagy induction enhances TDP43 turnover and survival in neuronal ALS models. Nat Chem Biol 10, 677–685, doi:10.1038/nchembio.1563 (2014).

22 Barmada, S. J. et al. Cytoplasmic mislocalization of TDP-43 is toxic to neurons and enhanced by a mutation associated with familial amyotrophic lateral sclerosis. J Neurosci 30, 639–649, doi:10.1523/jneurosci.4988-09.2010 (2010).

23 Jonsson, P. A. et al. Motor neuron disease in mice expressing the wild type-like D90A mutant superoxide dismutase-1. J Neuropathol Exp Neurol 65, 1126–1136, doi:10.1097/01.jnen.0000248545.36046.3c (2006).

24 Wainger, B. J. et al. Modeling pain in vitro using nociceptor neurons reprogrammed from fibroblasts. Nat Neurosci 18, 17–24, doi:10.1038/nn.3886 (2015).

25 Qin, H., Zhao, A., Ma, K. & Fu, X. Chemical conversion of human and mouse fibroblasts into motor neurons. Sci China Life Sci 61, 1151–1167, doi:10.1007/s11427-018-9359-8 (2018).

26 Steg, L. C. et al. Novel epigenetic clock for fetal brain development predicts prenatal age for cellular stem cell models and derived neurons. Mol Brain 14, 98, doi:10.1186/s13041-021-00810-w (2021).

27 Lapasset, L. et al. Rejuvenating senescent and centenarian human cells by reprogramming through the pluripotent state. Genes Dev 25, 2248–2253, doi:10.1101/gad.173922.111 (2011).

28 Wray, S. Modeling tau pathology in human stem cell derived neurons. Brain Pathol 27, 525–529, doi:10.1111/bpa.12521 (2017).

29 Tang, Y., Liu, M. L., Zang, T. & Zhang, C. L. Direct Reprogramming Rather than iPSC-Based Reprogramming Maintains Aging Hallmarks in Human Motor Neurons. Front Mol Neurosci 10, 359, doi:10.3389/fnmol.2017.00359 (2017).

30 Liu, M. L., Zang, T. & Zhang, C. L. Direct Lineage Reprogramming Reveals Disease-Specific Phenotypes of Motor Neurons from Human ALS Patients. Cell Rep 14, 115–128, doi:10.1016/j.celrep.2015.12.018 (2016).

31 He, K., Zhang, X., Ren, S. & Sun, J. in Proceedings of the IEEE conference on computer vision and pattern recognition. 770-778.

32 Huang, S. L. et al. A robust TDP-43 knock-in mouse model of ALS. Acta Neuropathol Commun 8, 3, doi:10.1186/s40478-020-0881-5 (2020).

33 Arrasate, M., Mitra, S., Schweitzer, E. S., Segal, M. R. & Finkbeiner, S. Inclusion body formation reduces levels of mutant huntingtin and the risk of neuronal death. Nature 431, 805–810, doi:10.1038/nature02998 (2004).

34 Martineau, É., Di Polo, A., Vande Velde, C. & Robitaille, R. Dynamic neuromuscular remodeling precedes motor-unit loss in a mouse model of ALS. Elife 7, doi:10.7554/eLife.41973 (2018).

35 Suk, T. R. & Rousseaux, M. W. C. The role of TDP-43 mislocalization in amyotrophic lateral sclerosis. Mol Neurodegener 15, 45, doi:10.1186/s13024-020-00397-1 (2020).

36 Arnold, E. S. et al. ALS-linked TDP-43 mutations produce aberrant RNA splicing and adult-onset motor neuron disease without aggregation or loss of nuclear TDP-43. Proc Natl Acad Sci U S A 110, E736–745, doi:10.1073/pnas.1222809110 (2013).

37 Ma, X. R. et al. TDP-43 represses cryptic exon inclusion in the FTD-ALS gene UNC13A. Nature 603, 124–130, doi:10.1038/s41586-022-04424-7 (2022).

38 Droppelmann, C. A., Campos-Melo, D., Moszczynski, A. J., Amzil, H. & Strong, M. J. TDP- 43 aggregation inside micronuclei reveals a potential mechanism for protein inclusion formation in ALS. Sci Rep 9, 19928, doi:10.1038/s41598-019-56483-y (2019).

39 Lanznaster, D., Hergesheimer, R., Vourc’h, P., Corcia, P. & Blasco, H. TDP43 aggregates: the ’Schrödinger’s cat’ in amyotrophic lateral sclerosis. Nat Rev Neurosci 22, 514, doi:10.1038/s41583-021-00477-1 (2021).

40 Schiff, L. et al. Integrating deep learning and unbiased automated high-content screening to identify complex disease signatures in human fibroblasts. Nat Commun 13, 1590, doi:10.1038/s41467-022-28423-4 (2022).

41 Bissaro, M. & Moro, S. Rethinking to riluzole mechanism of action: the molecular link among protein kinase CK1δ activity, TDP-43 phosphorylation, and amyotrophic lateral sclerosis pharmacological treatment. Neural Regen Res 14, 2083–2085, doi:10.4103/1673-5374.262578 (2019).

42 Fang, T. et al. Stage at which riluzole treatment prolongs survival in patients with amyotrophic lateral sclerosis: a retrospective analysis of data from a dose-ranging study. Lancet Neurol 17, 416–422, doi:10.1016/s1474-4422(18)30054-1 (2018).

43 Masuda, M. et al. TNIK inhibition abrogates colorectal cancer stemness. Nat Commun 7, 12586, doi:10.1038/ncomms12586 (2016).

44 Lee, Y., Jung, J. I., Park, K. Y., Kim, S. A. & Kim, J. Synergistic inhibition effect of TNIK inhibitor KY-05009 and receptor tyrosine kinase inhibitor dovitinib on IL-6-induced proliferation and Wnt signaling pathway in human multiple myeloma cells. Oncotarget 8, 41091–41101, doi:10.18632/oncotarget.17056 (2017).

45 Tu, P. H. et al. Transgenic mice carrying a human mutant superoxide dismutase transgene develop neuronal cytoskeletal pathology resembling human amyotrophic lateral sclerosis lesions. Proc Natl Acad Sci U S A 93, 3155–3160, doi:10.1073/pnas.93.7.3155 (1996).

46 Mullard, A. NfL makes regulatory debut as neurodegenerative disease biomarker. Nat Rev Drug Discov 22, 431–434, doi:10.1038/d41573-023-00083-z (2023).

47 Dawson, T. M., Golde, T. E. & Lagier-Tourenne, C. Animal models of neurodegenerative diseases. Nat Neurosci 21, 1370–1379, doi:10.1038/s41593-018-0236-8 (2018).

48 Mertens, J., Reid, D., Lau, S., Kim, Y. & Gage, F. H. Aging in a Dish: iPSC-Derived and Directly Induced Neurons for Studying Brain Aging and Age-Related Neurodegenerative Diseases. Annu Rev Genet 52, 271–293, doi:10.1146/annurev-genet-120417-031534 (2018).

49 Zhang, Y. et al. Prospects of Directly Reprogrammed Adult Human Neurons for Neurodegenerative Disease Modeling and Drug Discovery: iN vs. iPSCs Models. Front Neurosci 14, 546484, doi:10.3389/fnins.2020.546484 (2020).

50 Mendenhall, A. R., Martin, G. M., Kaeberlein, M. & Anderson, R. M. Cell-to-cell variation in gene expression and the aging process. Geroscience 43, 181–196, doi:10.1007/s11357-021-00339-9 (2021).

51 Sugano, T. et al. Pharmacological blockage of transforming growth factor-β signalling by a Traf2-and Nck-interacting kinase inhibitor, NCB-0846. Br J Cancer 124, 228–236, doi:10.1038/s41416-020-01162-3 (2021).

52 Sekita, T. et al. Feasibility of Targeting Traf2-and-Nck-Interacting Kinase in Synovial Sarcoma. Cancers (Basel) 12, doi:10.3390/cancers12051258 (2020).

53 Yang, Y. M. et al. A small molecule screen in stem-cell-derived motor neurons identifies a kinase inhibitor as a candidate therapeutic for ALS. Cell Stem Cell 12, 713–726, doi:10.1016/j.stem.2013.04.003 (2013).

54 Thams, S. et al. A Stem Cell-Based Screening Platform Identifies Compounds that Desensitize Motor Neurons to Endoplasmic Reticulum Stress. Mol Ther 27, 87–101, doi:10.1016/j.ymthe.2018.10.010 (2019).

55 Wu, C., Watts, M. E. & Rubin, L. L. MAP4K4 Activation Mediates Motor Neuron Degeneration in Amyotrophic Lateral Sclerosis. Cell Rep 26, 1143–1156.e1145, doi:10.1016/j.celrep.2019.01.019 (2019).

56 Bos, P. H. et al. Development of MAP4 Kinase Inhibitors as Motor Neuron-Protecting Agents. Cell Chem Biol 26, 1703–1715.e1737, doi:10.1016/j.chembiol.2019.10.005 (2019).

57 Delpire, E. The mammalian family of sterile 20p-like protein kinases. Pflugers Arch 458, 953–967, doi:10.1007/s00424-009-0674-y (2009).

58 Dan, I., Watanabe, N. M. & Kusumi, A. The Ste20 group kinases as regulators of MAP kinase cascades. Trends Cell Biol 11, 220–230, doi:10.1016/s0962-8924(01)01980-8 (2001).

59 Chuang, H. C., Wang, X. & Tan, T. H. MAP4K Family Kinases in Immunity and Inflammation. Adv Immunol 129, 277–314, doi:10.1016/bs.ai.2015.09.006 (2016).

60 Wang, Y. et al. Chemical conversion of mouse fibroblasts into functional dopaminergic neurons. Exp Cell Res 347, 283–292, doi:10.1016/j.yexcr.2016.07.026 (2016).

61 Tian, E. et al. Small-Molecule-Based Lineage Reprogramming Creates Functional Astrocytes. Cell Rep 16, 781–792, doi:10.1016/j.celrep.2016.06.042 (2016).

62 Cao, N. et al. Conversion of human fibroblasts into functional cardiomyocytes by small molecules. Science 352, 1216–1220, doi:10.1126/science.aaf1502 (2016).

63 Quek, H. et al. ALS monocyte-derived microglia-like cells reveal cytoplasmic TDP-43 accumulation, DNA damage, and cell-specific impairment of phagocytosis associated with disease progression. J Neuroinflammation 19, 58, doi:10.1186/s12974-022-02421-1 (2022).

64 Edgington, E. S. Randomization Tests. The Journal of Psychology 57, 445–449, doi:10.1080/00223980.1964.9916711 (1964).

65 Linsley, J. W. et al. Genetically encoded cell-death indicators (GEDI) to detect an early irreversible commitment to neurodegeneration. Nature Communications 12, 5284, doi:10.1038/s41467-021-25549-9 (2021).

